# ctsGRN: Inferring cell type-specific gene regulatory networks in *Arabidopsis*

**DOI:** 10.1101/2025.10.15.682501

**Authors:** Yu Jiang, Haolin Yang, Jian Gao, Qi Zhang, Zitai Yue, Xuemei Wei, Junpeng Zhang

## Abstract

Gene regulatory networks (GRNs) govern life processes. And exploring GRNs is of great value for understanding biological functions and developing species resources. At present, studies on plant GRNs are limited, mostly focusing on the tissue level, which hinders the discovery of cell type-specific regulatory networks and mechanisms. To address this, we have proposed a GRN research framework named ctsGRN (CellTypeSpecGRN) for conducting cell-specific GRN research at the cell type resolution. In this framework, we employed three representative algorithms from statistics (e.g., correlation analysis), traditional machine learning (e.g., random forest) and deep learning (e.g., neural networks) to construct GRNs for different cell types in *Arabidopsis* roots. Among these, traditional machine learning methods showed the best performance. Experimental validation through yeast one-hybrid (Y1H) assays and ChIP-seq data confirmed the high reliability and predictive power of these inferred GRNs. By analyzing these networks, we mined novel cell type-specific regulatory relationships, identified core transcription factor (TF) *AT5G08790* involved in root development, discovered 249 cell-type-specific hub TFs, and 136 key functional modules were found to be enriched in biological processes. Additionally, we revealed significant heterogeneity of TF-target gene interactions across various cell types. This comprehensive analysis offers a detailed overview of TF-mediated cell type-specific transcriptional regulation in *Arabidopsis* roots, providing new insights into the molecular mechanisms of root development and establishing a guideline framework for GRN research at the cell-type level in plants.

## Introduction

Accurate spatiotemporal gene transcription and expression are essential for life activities. Gene transcription is initiated when pioneer transcription factors (TFs) bind to DNA, triggering changes in chromatin accessibility. This process transforms DNA from a tightly compacted state to an open form, enabling other TFs to attach to distant cis-regulatory elements (CREs) on the DNA[1–3]. Cooperating with co-factors, these TFs collectively recruit and stabilize the RNA polymerase complex, which synthesizes mRNA using DNA as a template[4]. Various elements regulate this process, including TFs, splicing factors, long non-coding RNAs, microRNAs and metabolites, with TFs playing a critical role that can either enhance or repress the transcriptional rate of target genes[5, 6]. At the genomic level, the complex interactions among chromatin, TFs, and target genes form elaborate regulatory circuits defined as the gene regulatory networks (GRNs).

Recent advances in next-generation sequencing and computing technologies facilitated more accurate inference of GRNs using multi-omics data and mathematical approaches[7]. Therefore, GRNs serve as interpretable computational models that depict causal relationships and regulatory dynamics among genes by graphical or mathematical representation[8, 9]. Genome-scale datasets, including transcriptome, TFs binding sites information and chromatin accessibility etc., can be integrated into GRNs models to elucidate the regulatory mechanisms underlying complex biological processes[10, 11]. In GRN models, nodes represent genes, some of which are TFs, while the edges denote regulatory interactions between them. Hence, inferred GRNs can predict interactions between TFs and the genes they regulate, making GRNs an effective tool for identifying key regulatory and target genes involved in specific biological processes[12]. Moreover, uncovering the topology and dynamics of GRNs is fundamental to understand about how living cells are organized and operated, as well as how individual gene is regulated across different conditions, tissues or cell types[13].

Significant advancements have recently been achieved in inferring GRNs using both experimental and computational approaches. High-performing GRN inference algorithms have been continuously developed by using multi-omics data and other experimental datasets. Research on plant GRNs has also begun to gain momentum. By inferring GRN, researchers have uncovered new insights into the regulatory mechanisms governing gene expression across various biological contexts in plants. For example, four tissue-specific GRNs in maize were constructed utilizing GENIE3, a machine-learning algorithm applied to extensive RNA-Seq expression data, providing a standardized platform adaptable to any species with genome-wide expression data for GRN construction[14]. In soybeans, a robust unsupervised network inference scheme was used to predict GRNs related to the low phytic acid (lpa) trait, helping to elucidate molecular mechanisms affecting seed viability, germination and field emergence of crops[11]. GRNs inference algorithm MINI-EX identified key regulators involved in root development in *Arabidopsis thaliana* and rice, leaf development in *A.thaliana* and ear development in maize[15]. In addition, a network-based supervised learning approach for large-scale functional data integration was presented, resulting in an integrative GRN (iGRN) at tissue-level resolution in *Arabidopsis.* This iGRN demonstrated strong predictive power, accurately inferring known functions for TFs and predicting new functions for hundreds of previously uncharacterized TFs, including novel regulators of ROS signaling pathway[16]. Collectively, these research strategies and discoveries have accumulated valuable experience and knowledge for advancing plant GRN inference and analysis work.

Most plant GRN studies have been performed at the tissue level resolution, making it difficult to disentangle regulatory programs unique to specific cell types or states[17, 18]. At present, there is a lack of cell-type-specific GRNs derived from single-cell data in plants, which limits our understanding of the organization of specific cell types in plant. In this study, we constructed cell-type-specific GRNs for *Arabidopsis* roots at cell type resolution utilized three distinct algorithms. Machine learning algorithms demonstrates outstanding performance, and validation through Y1H assays and publicly available ChIP-seq data confirms the high predictive accuracy of the GRNs. Utilizing these high-quality networks, we identified numerous previously unknown regulatory relationships; We also pinpointed key TFs and functional modules specific to cell types which play important roles in different biological processes; Moreover, significant heterogeneity exists in TFs and their target genes across GRNs of different cell types. Our study provides a comprehensive overview of transcription factor-mediated GRNs in *Arabidopsis* root cells, offering valuable insights into the molecular mechanisms governing root development and establishing a reference framework for investigating plant biological processes at cellular resolution.

## Results

### Single-cell transcriptome of Arabidopsis roots reveals cell-type-specific gene expression patterns

Analyzing gene expression profiles at the cell-type level provides critical insights into the mechanisms of cell genesis and differentiation. In this study, single-cell transcriptomic data of *Arabidopsis* root from public database were examined to investigate gene expression characteristics related to root development. After data processing, a total of 22,017 genes were identified, and 3,018 high-quality *Arabidopsis* root cells were classified into nine distinct cell types, including Atrichoblast, Columella, Cortex, Endodermis, Lateral root cap, Metaphloem sieve element, Trichoblast, Mature and Xylem (Fig 1a). Among these, the Lateral root cap contains the largest number of cells, while the Mature cell type is the least represented (Fig 1b). Expression profile analysis reveals widespread activity of gene transcription across different cell types, with all except Trichoblast expressing over 15,000 genes. Trichoblast cells express the lowest genes (14,170 genes), whereas Metaphloem sieve element express the most (19,771 genes) (Fig 1c). Ubiquitous expression was observed in 10.08% of genes (2,219/22,017) across all nine cell types, whereas 4.25% (935/22,017) displayed cell type-specific expression patterns (Fig 1d). Analysis of the TFs expression profile shows that among the 2,296 TFs annotated in the PlantTFDB database, Metaphloem sieve element express the highest number, and Trichoblast the lowest, with most cell types expressing more than 1,000 TFs (Fig 1e); 29.27% (672/2,296) of TFs are expressed in all cell types, while 4.14% (95/2,296) show significant cell type specificity (Fig 1f). Further analysis reveals that highly expressed TFs (expression level > 1) are predominantly concentrated in Columella, Mature and Metaphloem sieve element cells, whereas lowly expressed TFs (expression level < 0.01) are found in Trichoblast and Lateral root cap (Fig 1g). The 672 TFs shared across all cell types exhibit diverse expression patterns. For instance, TFs highly express in Atrichoblast, Endodermis, Cortex, and Lateral root cap are expressed at low levels in other cell types. Notably, *AT5G08790* is highly expressed in all cell types, while *AT3G48440* and *AT5G23000* exhibit low expression universally. Some TFs, such as *AT3G44750*, *AT5G58010* show high expression specific to certain cell types, but low expression elsewhere; In Trichoblast cells, only *AT1G09540* exhibits relatively high expression. These findings demonstrate that different cell types possess unique expression profiles for both non-TFs and TFs, providing valuable insights into the cell type-specific transcriptional regulatory mechanisms that govern root development in *Arabidopsis*.

**Fig 1.**
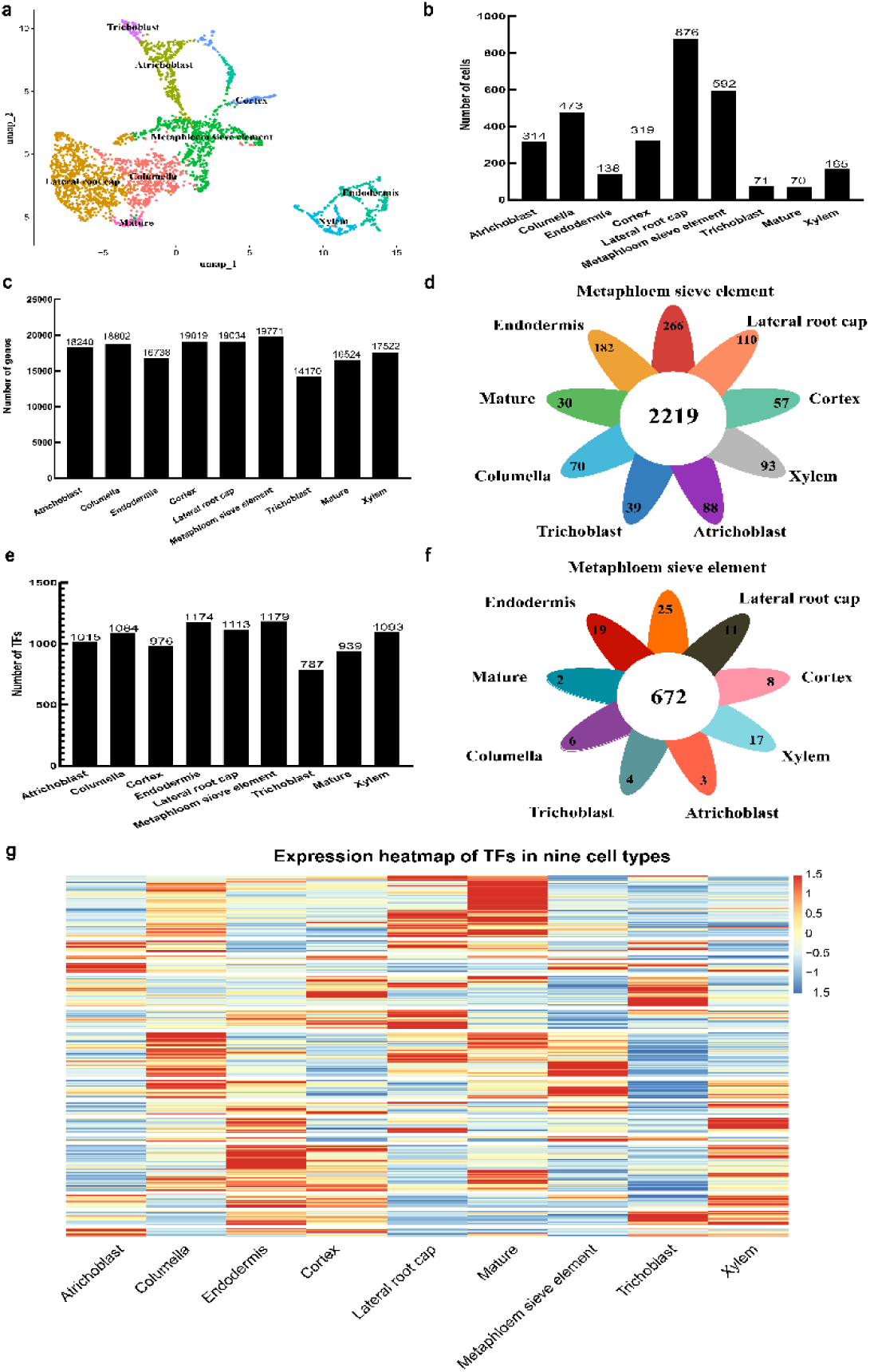
The expression profiles analysis of genes across cell-type in *Arabidopsis* root. (a) A UMAP map showing the number of single cells from *Arabidopsis*; (b) Number of cells in each cell type; (c) A histogram displaying the number of genes expressed within the nine cell types; (d) A distribution map showing the number of genes specifically expressed in each cell type. Color represents genes specifically expressed in each cell, and white represents genes expressed in all cells; (e) Number of TFs in nine different cell types; (f) The number of TFs specifically expressed in each cell type. Color represents the TFs specific to each cell type, and white represents TFs expressed in all cell types; (g) Expression heat map of 672 TFs that are expressed in all nine cell types.

### Construction of cell-specific gene regulatory networks in A.thaliana root

GRN inference algorithms are primarily designed based on concepts from statistical methods, traditional machine learning and deep learning. Accordingly, we employed three representative algorithms embodying these principles—WGCNA, GENIE3 and DeepSEM—to infer cell-type-specific GRNs in the *Arabidopsis* root. The results indicate that the data outputs from these three algorithms differ across cell types. WGCNA produces the largest number of outputs, followed by DeepSEM with a moderate amount, while GENIE3 generates the fewest (Table 1).

**Table 1.** Input data and network construction output data table.

| Cell Type | Cell count | Gene type |  | Regulatory relationship count |  |  |
| --- | --- | --- | --- | --- | --- | --- |
|  |  | Gene | TF | WGCNA | GENIE3 | DeepSEM |
| Atrichoblast | 314 | 18240 | 1015 | 1926658 | 671640 | 994173 |
| Columella | 473 | 18802 | 1084 | 1627689 | 714335 | 1004346 |
| Cortex | 138 | 16738 | 976 | 1894810 | 611199 | 778609 |
| Endodermis | 319 | 19019 | 1174 | 2752913 | 780066 | 1199616 |
| Lateral root cap | 876 | 19034 | 1113 | 2019000 | 739324 | 972598 |
| Metaphloem sieve element | 592 | 19771 | 1179 | 2954829 | 802006 | 1139320 |
| Trichoblast | 71 | 14170 | 787 | 1054969 | 440003 | 549511 |
| Mature | 70 | 16524 | 939 | 952815 | 598344 | 776765 |
| Xylem | 165 | 17522 | 1093 | 1978281 | 694387 | 1022872 |

AUPR(Area Under the Precision-Recall Curve) was caculated using published ChIP-seq, ATAC-seq, and DNase-seq datasets to evaluate the performance of the three algorithms for network inference. The results show that GENIE3 achieves an AUPR value of 0.865, slightly outperforming WGCNA and DeepSEM (Fig 2a), confirming its status as one of the leading methods for reconstructing GRNs from scRNA-seq data, consistent with previous findings[19]. In terms of computational efficiency, WGCNA has the shortest runtime, followed by GENIE3, while DeepSEM exhibits the highest computational complexity and has the longest runtime due to its deep learning framework (Fig 2b). The network topology analysis revealed that the GRNs inferred by all three algorithms exhibited typical scale-free and small-world network characteristics (S1 Fig). Their node degree distributions follow a power-law pattern, and network density is significantly higher than those of comparable random networks, whereas characteristic path lengths show minimal significant differences from random network. Notably, networks constructed by DeepSEM exhibit significantly higher density compared with those generated by WGCNA and GENIE3, suggesting a more densely connected topology (S1 Fig).

**Fig 2.**
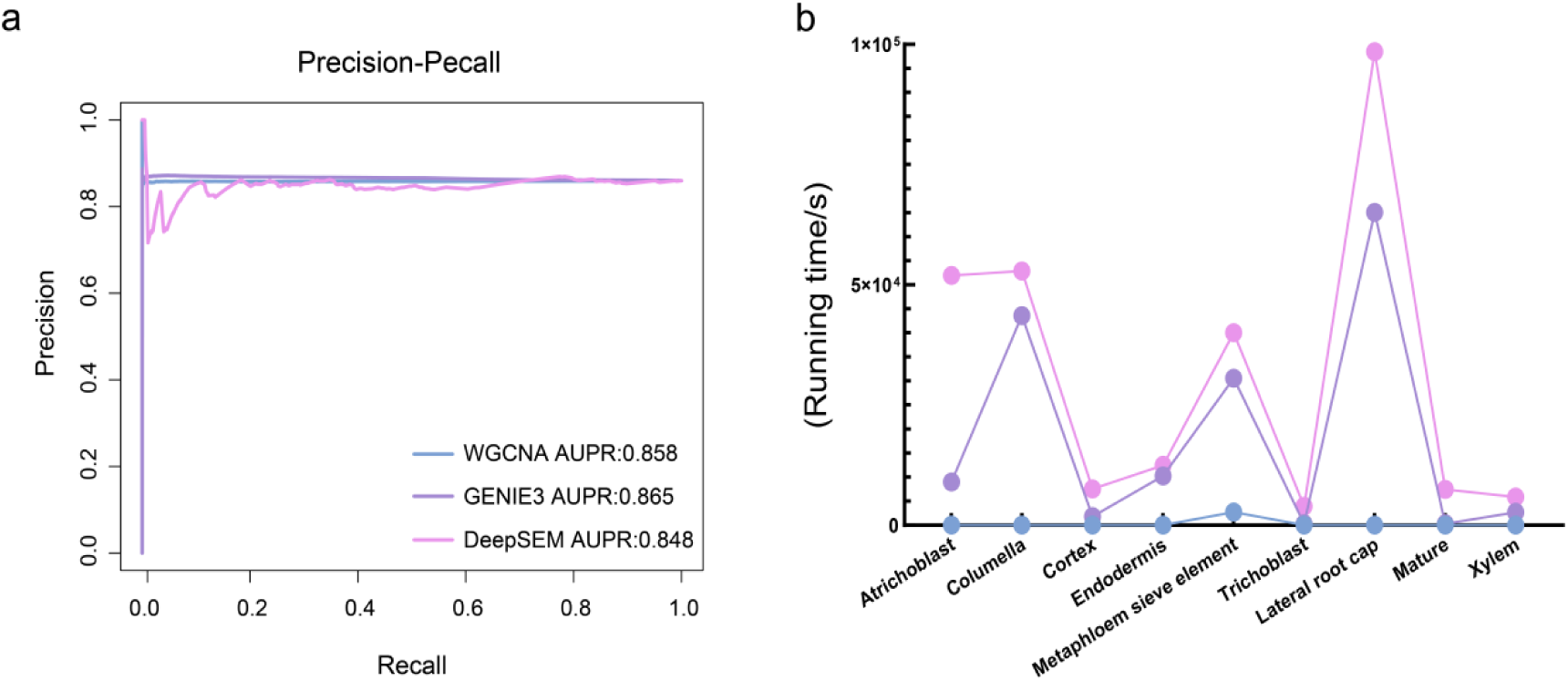
The performance of three algorithms for GRN construction. (a) Assessment for algorithms based on AUPR; (b) The running time taken by the algorithms.

According to the evaluation results presented above, GENIE3 shows superior performance, while DeepSEM performs adequately in certain situations but requires further optimization. WGCNA shows a low validation rate and is recommended only for specific purpose or when used alongside other validation methods. Consequently, the GRN constructed by the machine learning algorithm GENIE3 will be selected for further analysis of prediction performance, network structure, key regulatory factors, and regulatory function analysis.

### GRNs demonstrate prediction performance for cell-type-specific regulatory interactions

If GRN predictions are enriched for ChIP-seq confirmed regulatory relationships, which would suggest that GRN could reliably identify putative regulatory relationships for other TFs without ChIP-seq data[20]. Accordingly, we compared the numbers of regulatory interactions predicted by GRNs across nine cell types and assessed their overlap with interactions validated by ChIP-seq (Fig 3, S2 Table). Hypergeometric distribution test was applied to determine the significance of the overlapping regulatory interactions. The results indicate that most GRN-predicted regulatory pairs are significantly enriched among ChIP-identified regulatory targets (*p* < 0.05), suggesting that the GENIE3 inferred networks can effectively capture authentic biological regulatory interactions and uncover novel previously uncharacterized regulatory relationships. Utilizing this network, we predicted a total of 1,176,264 novel regulatory pairs across nine cell-type-specific regulatory networks, comprising 2,225 pairs unique to specific cell types and 1,174,039 pairs common across cell types (S3 Table).

**Fig 3.**
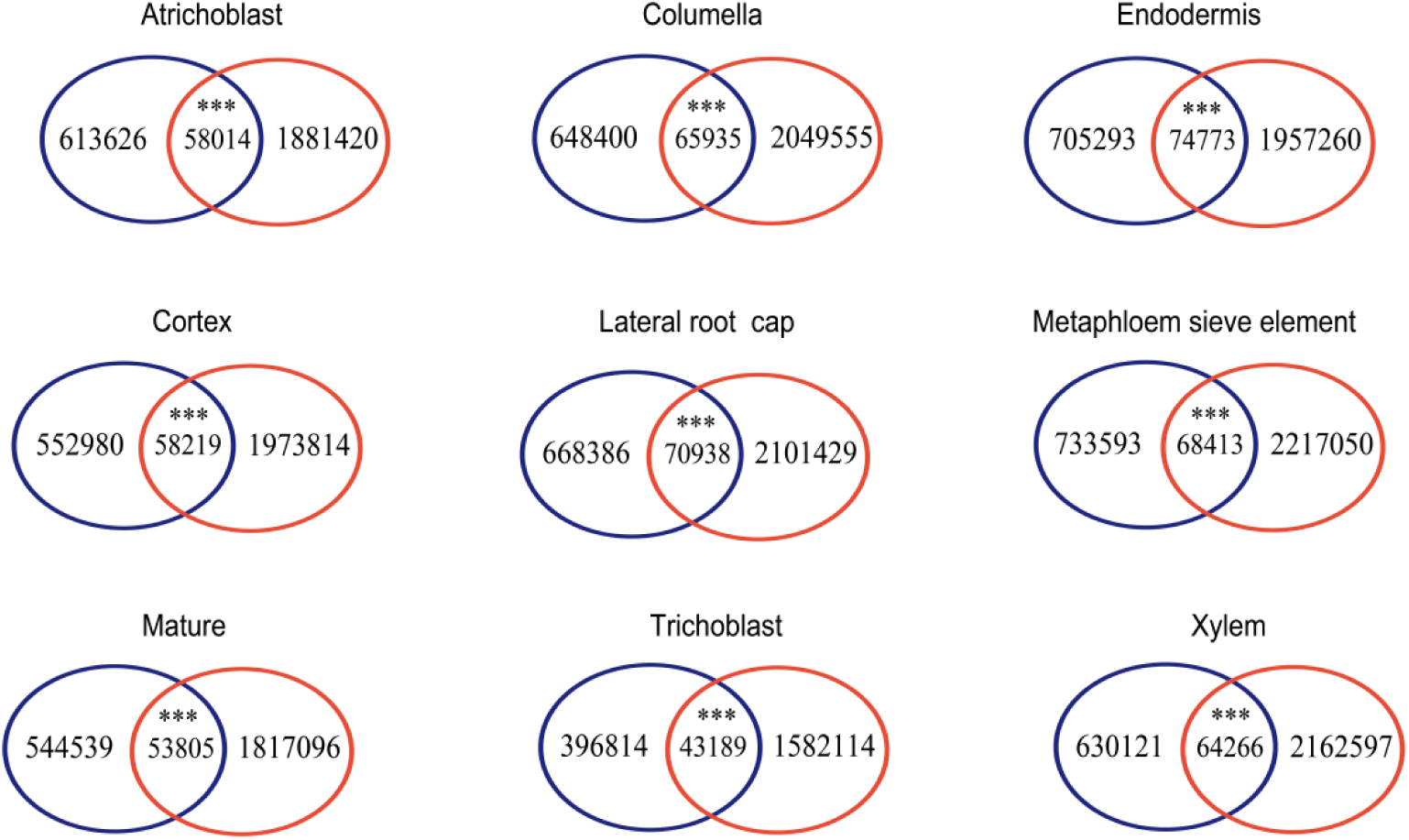
Venn diagram summarizing the overlap between predicted regulatory relationships and ChIP-seq. Blue circles are the number of predicted regulatory relationships for cell type-specific GRNs; red circles are the number of regulatory relationships identified by ChIP-seq. p-values were calculated using the hypergeometric distribution test, and *** *(p-value* less than 0.01) or * (*p-value* less than 0.05) indicate significant overlap.

### Yeast one-hybrid experiments confirm that the transcription factor regulatory relationship predicted by GRN is real

To validate whether the regulatory relationships between key TFs and target genes predicted by GRN model truly exist, we randomly selected a pair consisting of a key TF and a target gene - *AT1G62300* (*WRKY6*) and *AT4G22710* (*CYP706A2*) - which have high-confidence predicted regulatory relationship, for experimental validation using the Y1H assay. Sequence analysis showed that one binding site of *WRKY6* was located in the promoter region of *CYP706A2* (from 18 to 36bp upstream of ATG). In the Y1H assay, the fragments of the *CYP706A2* promoter containing the binding site of WRKY6 were used as bait, all the yeast cells grew normally on the SD/-Leu medium without 50 ng/mL AbA. However, when 50 ng/mL AbA was added, growth continued in both the positive control and bait-prey, whereas the AD-CYP706A2 were completely inhibited (Fig 4). These results indicate that WRKY6 interacts with the promoter fragment in yeast, confirming that *CYP706A2* as a potential target gene of *WRKY6*, demonstrating that the outstanding predictive performance of our constructed GRN model.

**Fig 4.**
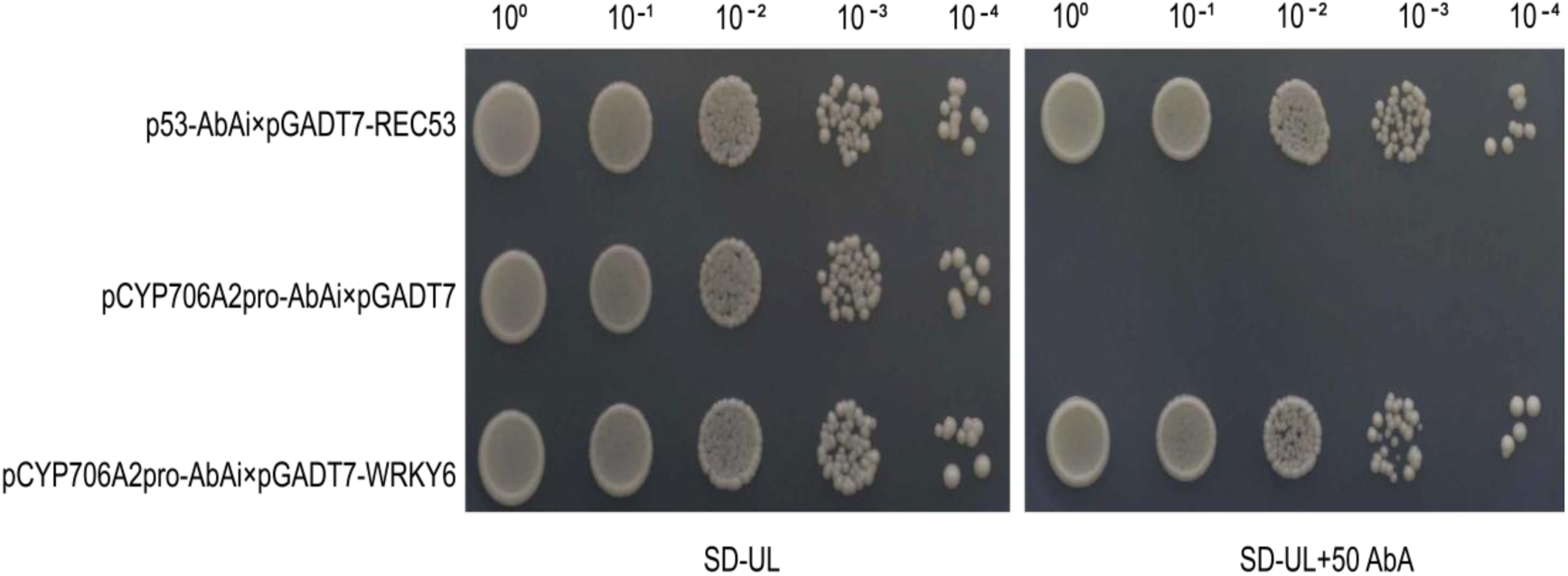
Y1H Gold yeast cells of bait-prey co-transformation on SD/-Leu medium with or without 50 ng/mL AbA.

### Network structure analysis reveals the complexity of gene expression regulation

Systematic analysis of the structural characteristics of inferring GRNs reveals that each gene is regulated by an average of 227 TFs, with approximately 49.91% of genes being regulated by more than this average (S4 Table). Conversely, each TF controls an average of 3646 target genes. In addition, 56% of TFs control more than 3,000 target genes, while 4.2% of genes are regulated by 50 to 100 TFs (S4 Table). It is worth noting that the number of regulatory edges in the GRNs varies among cell types; for example, the Metaphloem sieve element contains 802,006 edges, whereas the Trichoblast has only 440,003 edges. The average number of TF regulatory edges per TF ranges from 559 to 680 (Table 2), reflecting the cell-type-specific heterogeneity in network scale and regulatory intensity. The TF with the highest number of regulatory edges (*AT3G44750*), which is functionally annotated in the QuickGO database(https://www.ebi.ac.uk/QuickGO/), primarily mediates developmental polarity establishment and epigenetic regulation (Table 2). Conversely, the TF with the fewest regulatory edges (*AT5G23000*), mainly governs transcription and cell fate determination (Table 2). The core TF *AT5G08790* exhibits ubiquitous high expression and pivotal regulatory functions across multiple cell types (Table 2, Fig 1g), establishing its central role in the gene expression regulatory network of *Arabidopsis* root cells, suggesting a strong link between its high expression and gene transcription regulation. Comparison of the degree centrality of the shared TFs across different cell types reveals significant functional divergence in their regulatory roles (Fig 5a) and exhibiting striking cell-type-specific regulatory patterns. The coefficient of variation (CV) for degree centrality of the shared TFs across nine cell types exhibits a wide range, from 17.58 to 125.14, with an average of 42.48. The TF with the highest variability is *AT4G29100* (CV = 125.14) in the Endodermis. Among the top 100 TFs with the highest CV 27 are identified in the Endodermis, 18 in the Lateral root cap, and 17 in the Metaphloem sieve element, each exhibiting high degree centrality in their respective cell types (Fig 5b, S5 Table). Notably, some TFs show almost no regulatory edges in certain cell types, indicating minimal predicted target genes and limited regulatory activity in those cellular contexts. Collectively, these TFs form highly complex regulatory networks, which constitute the molecular regulatory basis for root cell differentiation and development in *Arabidopsis*.

**Fig 5.**
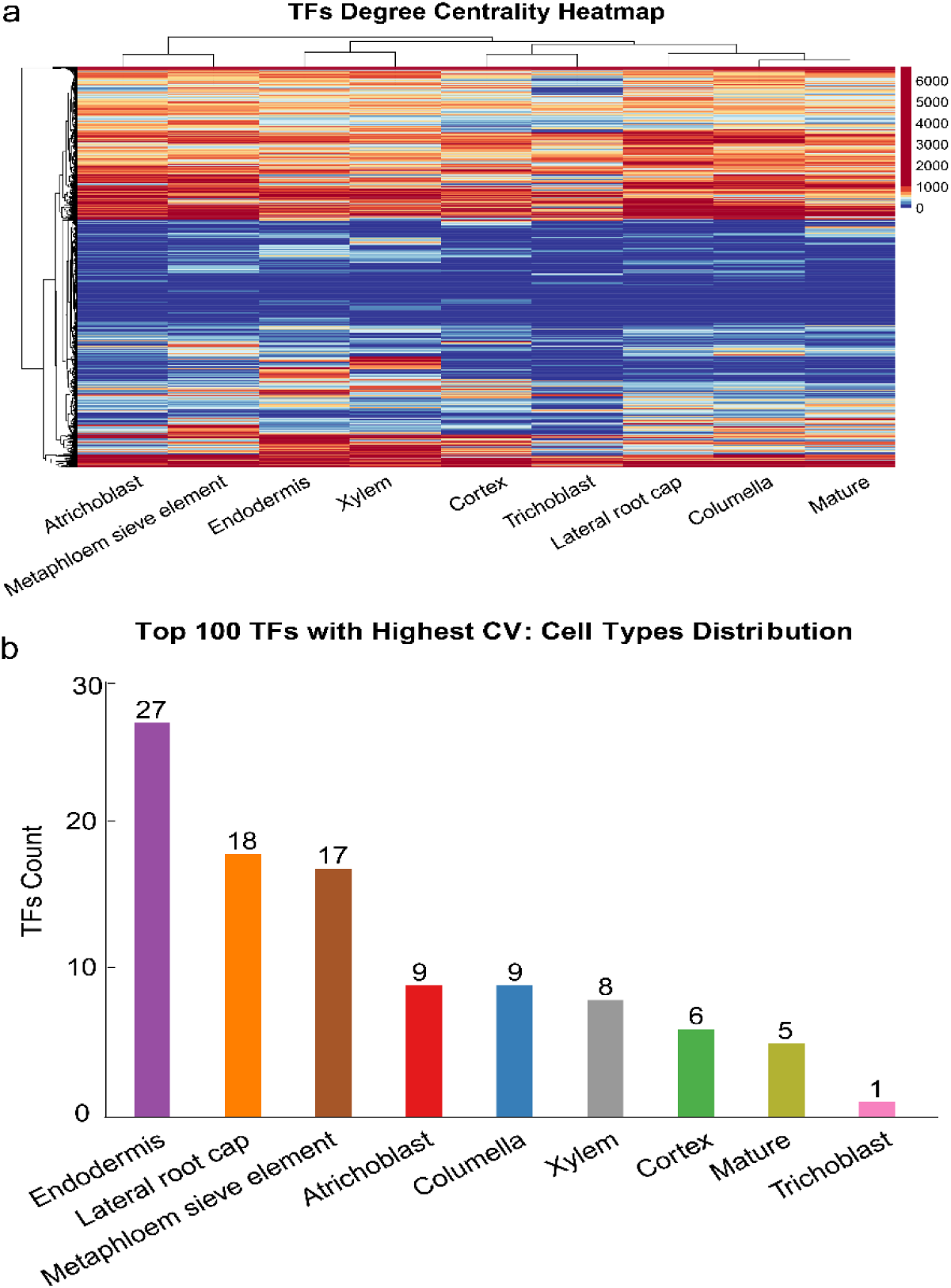
The variation of TF-mediated interacts in nine cell type-specific GRNs. (a) Heatmap showing the number of target genes regulated by TFs across nine cell-type-specific GRNs. The color scale is based on quantile segmentation, with each color representing 10% of the data. Hierarchical clustering is based on Euclidean distance; (b) The number of TFs with high coefficient of variation in nine different cell types.

**Table 2.** Scale and strength of TFs regulatory network in different cell types.

| Cell Type | Regulatory relationship | Maximum number of edges<br>TF (Number of edges) | Minimum number of edges<br>TF (number of edges) | Average number of<br>regulated edges |
| --- | --- | --- | --- | --- |
| Atrichoblast | 671640 | AT3G09735<br>(4884) | AT5G23000<br>(21) | 661.71 |
| Columella | 714335 | AT5G08790<br>(4825) | AT1G75250(55) | 658.98 |
| Cortex | 611199 | AT3G01220<br>(3430) | AT1G09710(47)<br>AT4G14540(47) | 626.23 |
| Endodermis | 780066 | AT5G08790<br>(4182) | AT5G11590<br>(50) | 664.45 |
| Lateral root cap | 739324 | AT5G08790<br>(5502) | AT1G35515(45)<br>AT5G64060(45) | 664.26 |
| Metaphloem<br>sieve element | 802006 | AT5G08790<br>(2910) | AT5G18090<br>(106) | 637.21 |
| Trichoblast | 440003 | AT5G63790<br>(2541) | AT1G31140<br>(66) | 559.09 |
| Mature | 598344 | AT3G44750<br>(6639) | AT1G26610<br>(30) | 680.24 |
| Xylem | 694387 | AT2G37120<br>(3136) | AT3G04450<br>(70) | 635.30 |

### Analysis of key transcription factors in the inferring GRNs reveals heterogeneity in cellular gene expression regulation

TFs typically play dominant roles in gene expression regulation. Thus, to identify key TFs that hold dominant regulatory influence, we applied a stringent threshold based on degree centrality. TFs with a degree centrality greater than 2,000 in any of the nine GRNs were considered key regulators. Using this criterion, we identified a total of 249 unique key TFs across all networks (Fig 6a, S2 Fig, S6 Table). Among these, the Lateral root cap contains the largest number of key TFs (47), while the Trichoblast has the lowest (4 key TFs). Overlap analysis shows that 27.71% (69/249) of these key TFs are exclusive to a single cell type, whereas two key TFs *AT5G08790* and *AT5G63790* are common to all nine cell types (Fig 6b). Previous studies indicate that both participate in DNA-templated transcriptional regulation and the development of multicellular organisms[21, 22]. Moreover, *AT5G08790* exhibits diverse functions, responding to various external stimuli such as light stimulation, wounding, fungal infection, sucrose, salicylic acid and jasmonic acid, thereby regulating multiple biological processes[23]. This is consistent with the previous findings demonstrating the significant regulatory role and core functional position of *AT5G08790* across *Arabidopsis* root cell types (Table 2). Further analysis reveals significant differences in the regulatory relationships of *AT5G08790* and *AT5G63790* among cell types (Fig 6c). Specifically, they share 150 and 101 common target genes, respectively, while possessing 4,088 and 2,438 unique target genes, respectively (Fig 6d). Correlation analysis of key TFs shows substantial regulatory heterogeneity across different cell types (Fig 7e). The greatest heterogeneity is observed between the Columella and Endodermis (correlation = 0.26), whereas the least heterogeneity is found between the Trichoblast and Metaphloem sieve element (correlation = 0.86), reflecting the dynamic variability of the regulatory network during the differentiation-to-maturation process in *Arabidopsis* root cells.

**Fig 6.**
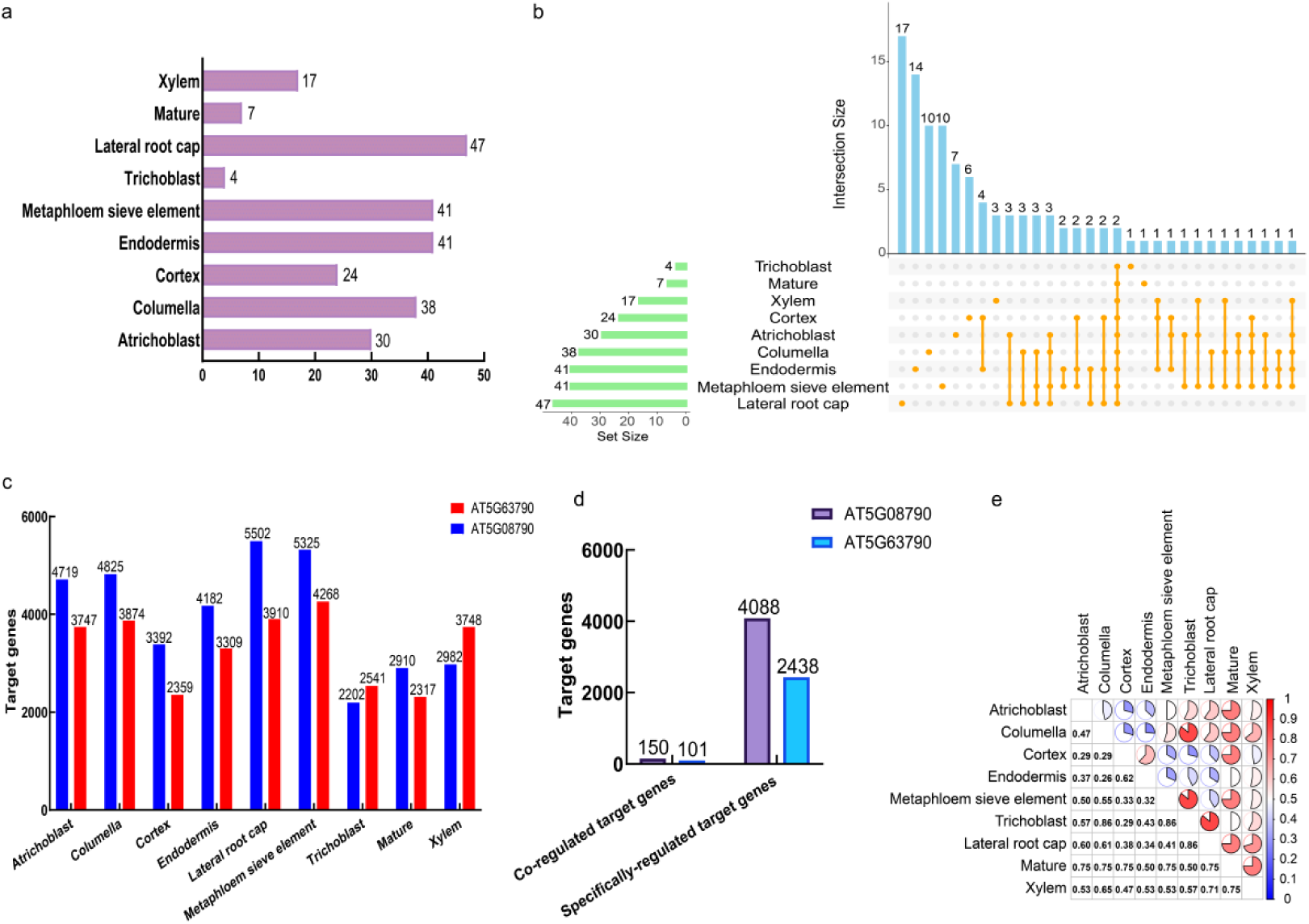
Identification and analysis of key TFs in nine cell type-specific GRNs. (a) The number of key TFs identified in the nine different cell types; (b) An Upset graph visualizing the set interactions among key TFs in each cell type GRN. Number of key TFs expressed in each individual or combination of cell types were represented in bar plot (blue). The intersections were represented by connected orange dots. Total number of key TFs expressed in each cell types were represented in green bar plot; (c) The number of regulatory relationships of *AT5G63790* and *AT5G08790* the nine different cell types; (d) The number of genes specifically regulated and co-regulated by *AT5G63790* and *AT5G08790* in the nine cell types; (e) Correlation analysis of key TFs in each cell type.

**Fig 7.**
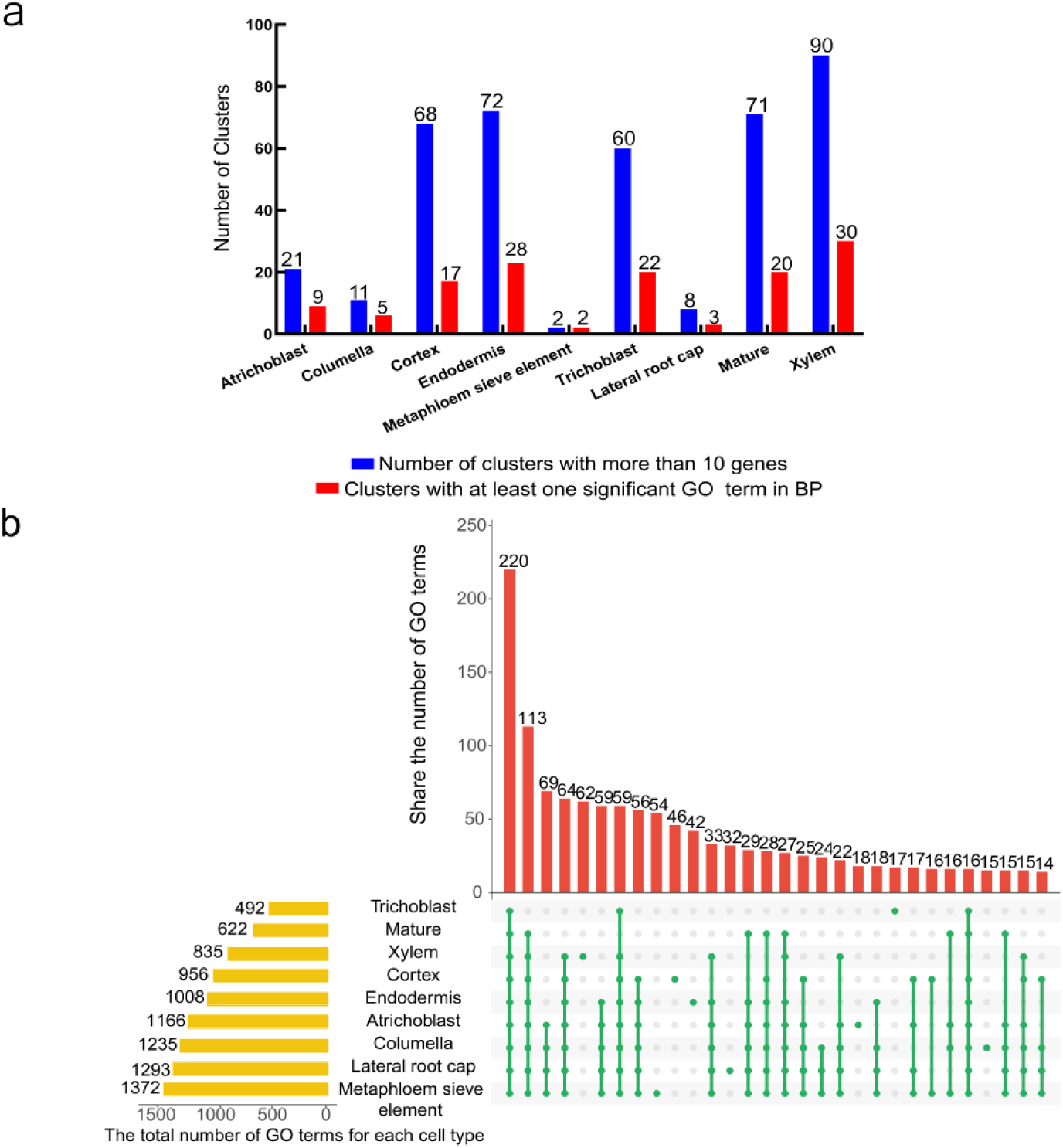
Identification of functional modules in cell-type-specific GRNs. (a) Number of functional modules in each GRNs; (b) Upset graph visualizes the distribution of enriched pathways in each GRNs. Red bars represent the number of enriched pathways in different cell types (red). The intersections were represented by connected green dots. Yellow bars represent the total number of enriched pathways in each cell type.

### Homology comparison and analysis reveals the functional characteristics of key TFs

*A.thaliana* and *Brassica napus* both belong to the *Brassicaceae* family and share an ancient whole-genome triplication (WGT) event. Approximately 75% of the genes in *B.napus* can be mapped to *A.thaliana* orthologs through microcolinearity blocks, which serve as evolutionary reference points for functional studies[24]. In this study, we integrated the genomic data of *B.napus* to conduct the phylogenetic and homology analysis on 10 previously unverified key TFs in *A. thaliana*, aiming to predict their biological functions. The results show that these 10 TFs exhibit strong homology with gene families in *B. napus* including HD-ZIP, LBD, NAC, ERF, among others (Table 3, S3 Fig).

**Table 3.** Homologous genes of key TFs in *Arabidopsis*.

| Key TFs | Homologous gene in *B.napus* | Homologous family |
| --- | --- | --- |
| AT1G10170 | BnaC08g14030D; BnaC08g42350D; BnaA08g25950D;<br>BnaA09g48270D; BnaC05g07520D; BnaA06g05950D; | NF-X1 |
| AT1G26960 | BnaC08g48560D; BnaC07g12560D; BnaA08g20030D; | HD-ZIP |
| AT1G74500 | BnaAnng30830D; BnaA07g22200D; BnaA07g31610D; | bHLH |
| AT3G23690 | BnaC06g35430D; BnaC06g22940D; BnaC06g22930D;<br>BnaA01g23600D; BnaCnng29920D; BnaA01g36300D;<br>BnaA07g06260D; BnaC07g07840D; |  |
| AT2G30130 | BnaC03g16640D; BnaA03g56340D; BnaA05g12220D;<br>BnaC04g54590D; | LBD |
| AT2G37120 | BnaCnng30990D; BnaAnng24190D; BnaC06g14570D;<br>BnaA09g33670D; | S1Fa-like |
| AT2G45050 | BnaC04g03950D; BnaC04g50080D; BnaA04g26010D;<br>BnaA05g04530D | GATA |
| AT3G29035 | BnaA09g02500D; BnaCnng49350D; BnaC07g25530D;<br>BnaA06g31080D; | NAC |
| AT4G17490 | BnaA01g34910D ; BnaC01g10080D;BnaCnng05080D ; | ERF |
| AT4G17500 | BnaAnng21280D; BnaC01g10100D; BnaA08g08310D |  |

Experimental validations such as quantitative PCR (qPCR), subcellular localization and yeast two-hybrid assays have demonstrated that HD-ZIP in *B.napus* interacts with HB12 homologs (BnaA.HB12.a, BnaC.HB12.a, BnaC.HB12.b) to form homodimers in response to abiotic stress[25]. The LBD family is speculated to participate in hormone-mediated stress regulation pathways (including root development, seed maturation, etc.) [26]; NAC family genes respond to abiotic stress and hormone signals, exhibit specific subcellular localization-related functions and can induce cell death[27]. The ERF family regulates cold stress signaling pathways, responds to various hormone signals, and dynamically controls the timing of cold adaptation process[28]. These results suggest that the *A.thaliana* genes *AT1G26960*, *AT2G30130*, *AT3G29035*, *AT4G17490* and *AT4G17500*, which are homologous to the aforementioned *Brassica* genes, likely share similar potential functions and participate in regulating specific biological processes.

### Identification and analysis of network functional modules reveal functional heterogeneity in Arabidopsis root cells

Functional module analysis aims to elucidate the mechanism behind life processes. To this end, we employed the MCL (Markov Cluster) algorithm to identify the functional modules within the inferring GRNs of nine different cell types and performed multi-level functional enrichment analysis (Fig 7, S7 Table). Our findings show significant differences in the number of functional modules across cell types (Fig 7a), with 33.75% of these modules (136/403) containing at least one significantly enriched biological process GO term (*adj.p* < 0.05). The upset graph illustrates both conserved and cell type-specific regulatory patterns among the enriched pathways in each module, of which there are 15 to 62 unique pathways and 14 to 220 conserved enriched pathways (Fig 7b). The core shared pathways mainly involve fundamental biological processes such as RNA splicing (present in 7 of 9 cell types) and ATP metabolism (6 of 9) (Fig 8a-b). In contrast, cell type-specific pathways include “glucuronoxylan biosynthesis” unique to Xylem, “asymmetric cell division regulation” specific to Trichoblast, and “sieve tube development” exclusive to Metaphloem sieve element (Fig 8a-b). Functional similarity analysis shows that Columella shares the highest functional similarity with Lateral root cap (Jaccard index = 0.8), while Metaphloem sieve element exhibits significant functional uniqueness (Fig 8c). Further analysis demonstrates that all cell types display significant functional heterogeneity but retain fundamental metabolic pathways (such as ribosome biogenesis, protein folding, and cellular respiration) (Fig 8d). Specifically, Xylem is enriched in pathways related to secondary cell wall biosynthesis and metabolism, aligning with its role in structural support; Trichoblast shows enrichment in oxidative stress response and protein transport pathways, reflecting its adaptation to the soil environment, and Lateral root cap is enriched in wound response and Golgi vesicle transport pathways, corresponding to its protective function. These results reveal that *Arabidopsis* root cells preserve essential physiological functions through conserved core pathways, while achieving functional differentiation with the help of cell type-specific pathways, establishing a model of “conserved basic functions-specialized functional differentiation”. These findings provide valuable insights into the complex regulatory networks that govern plant organ development.

**Fig 8.**
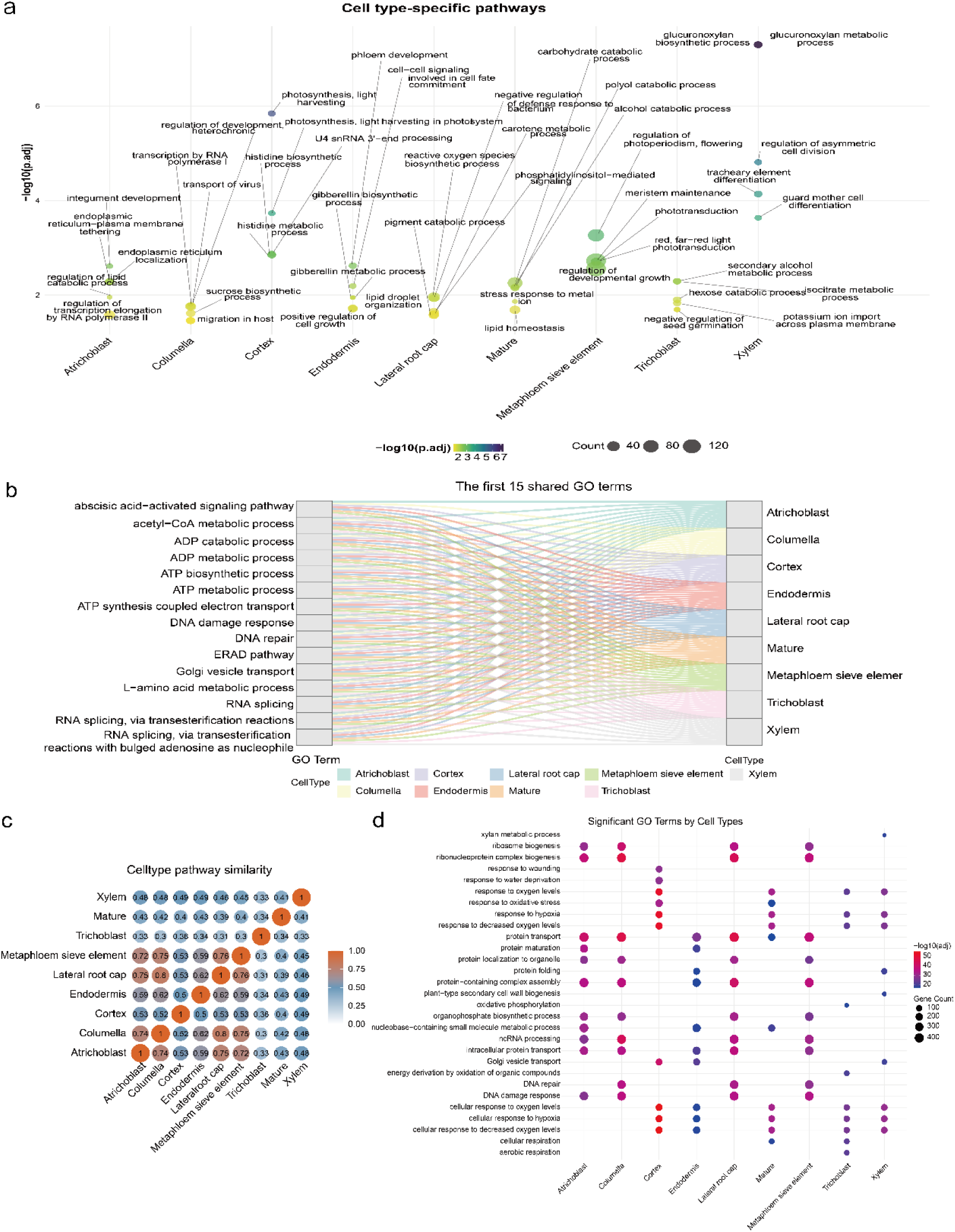
Analysis of functional modules in cell-type-specific GRNs. (a) Bubble diagram displaying the core enriched pathways specific to each cell type; (b) Sankey diagram showing the core enriched pathways shared by each cell type; (c) Similarity analysis of enriched pathways for each cell type; (d) Bubble diagram showing core enriched pathways for each cell type.

## Discussion

Research plant GRNs is essential for understanding biological processes and developing genetic resources. However, there are limited reports focusing on plant cell-type-specific GRNs utilizing the comprehensive inference strategies. In this study, we integrated single-cell transcriptomic data from *Arabidopsis* root with multiple GRNs inference algorithms to construct a high-quality, cell-type-specific root GRN with predictive performance. We evaluated the performance of three representative algorithms — WGCNA, GENIE3, and DeepSEM — based on statistical method, traditional machine learning, and deep learning approaches, respectively, using test data. Among them, GENIE3 offers the optimal balance between AUPR of 0.865 and computational time of 5.05h. In terms of parameter optimization, we set nTree to 1000 to maintain prediction accuracy while avoiding overfitting. This choice both addresses the signal loss caused by conservative parameters (nTree = 500) in a small gene set (<1,000) and avoids the computational burden from high settings (nTree = 5000) in large-scale datasets[29], providing a useful guideline for plant single-cell GRN inferring. For evaluating the predictive performance of GRNs, the network inferred by GENIE3 was validated by significant enrichment in ChIP-seq analysis (*p* < 0.05) and experimentally confirmed novel regulatory relationship via yeast one-hybrid assay.

Compared with previous root cell differentiation model[30], we systematically uncovered the heterogeneity characteristics of *Arabidopsis* root cell GRNs through cell-type resolution network analysis. We found highly intricate regulatory structure: on average, each gene is regulated by 227 TFs, while core TFs (56%) could regulate more than 3,000 target genes. Cell type-specific GRN analysis shows significant differences in network size (ranging from 440,003 to 802,006 edges), among which core TF (*AT5G08790*) play a key regulatory role across multiple cell types, including Columella and Endodermis, where it is involved in transcriptional regulation, development, and environmental responses[31]. Network structure analysis reveals that unique regulatory characteristic among cell types, with the Endodermis containing the highest number of highly variable TFs. We measured the regulatory heterogeneity across cell types (CV=17.58-125.14) and notably identified that 27.71% of key TFs are specific to cell types, a proportion significantly higher than previous estimates derived from bulk RNA-seq data[32].

The identification of core TFs like *AT5G08790* not only verified the hypothesis regarding pleiotropic TFs[33] but also improved the prediction accuracy to experimentally validated levels through yeast single-hybrid assays, thereby addressing the lack of experimental validation noted in earlier studies[34]. In addition, the ubiquitous high expression of this TF across all cell types suggests its potentially important role in the development of *Arabidopsis* roots. This finding is consistent with the previous research showing that *AT5G08790* (*ATAF2*) coordinates developmental processes in roots and other tissues via the JA signaling pathway[32]. Functional module analysis identified 26 core modules, with Module 2 in the Columella, associated with gibberellin signaling, closely matching previously described gibberellin regulatory networks. Additionally, the xylem-specific metabolism module broadens our understanding of xylem development[35, 36].

In this study, we conducted a comprehensive analysis of the evolutionary conservation and functional heterogeneity of key TFs in Arabidopsis root cells by integrating phylogenetic analysis and network functional module identification. Homology comparisons show that 10 candidate key TFs have highly conserved orthologs in *B.napus*, with 7 of these expressed across multiple cell types of Arabidopsis roots. Crucially, their DNA binding domains are highly conserved, indicating functional preservation in the Brassicaceae. Functional module analysis reveals a regulatory pattern characterized by “basic functional conservation-specialized functional differentiation“: all cell types maintain core metabolic pathways such as RNA splicing and ATP metabolism), yet display distinct functional heterogeneity—Xylem cells specifically enrich secondary cell wall synthesis pathways, Trichoblast uniquely participates in asymmetric cell division regulation, whereas Metaphloem sieve element specializes in the sieve tube development pathway—aligning closely with their biological roles. Notably, Columella and Lateral root cap show the highest functional similarity, whereas Metaphloem sieve element exhibits significant functional uniqueness. Furthermoure, we identified 26 core functional modules, including those related to hormone signaling and environmental responses. These findings not only elucidate the molecular regulatory networks of plant root cell differentiation but also provide potential target genes for genetic improvement of crop roots systems, especially conserved key TFs and their mediated functional modules in Brassicaceae crops.

Building upon this study, future studies could: 1) Integrate single-cell multi-omics data to construct GRNs; 2) Apply temporal network analysis methods to delineate the dynamic regulatory mechanisms involved in root development; 3) Leverage CRISPR-Cas9 gene editing technique to verify the functions of key TFs. These advancements will facilitate further refinement of GRN performance, provide a comprehensive regulatory map of plant root development and offer a molecular blueprint for designing root systems.

## Materials and Methods

### Research framework for this study

This study focused on the transcriptomic data from different cell types in the roots of the model plant *A.thaliana* and constructed cell type-specific GRNs for *Arabidopsis* roots by integrating three representative algorithms based on statistical, traditional machine learning, and deep learning approaches. The performance of these algorithms and the quality of the reconstructed GRNs were then assessed using external datasets. Finally, bioinformatic approaches were used to systematically map the landscape of cell type-specific GRNs, identify key regulators and functional modules within the network, predict novel regulatory interactions, and uncover the molecular mechanisms underlying root growth and development in *Arabidopsis* at cell-type resolution (Fig 9).

**Fig 9.**
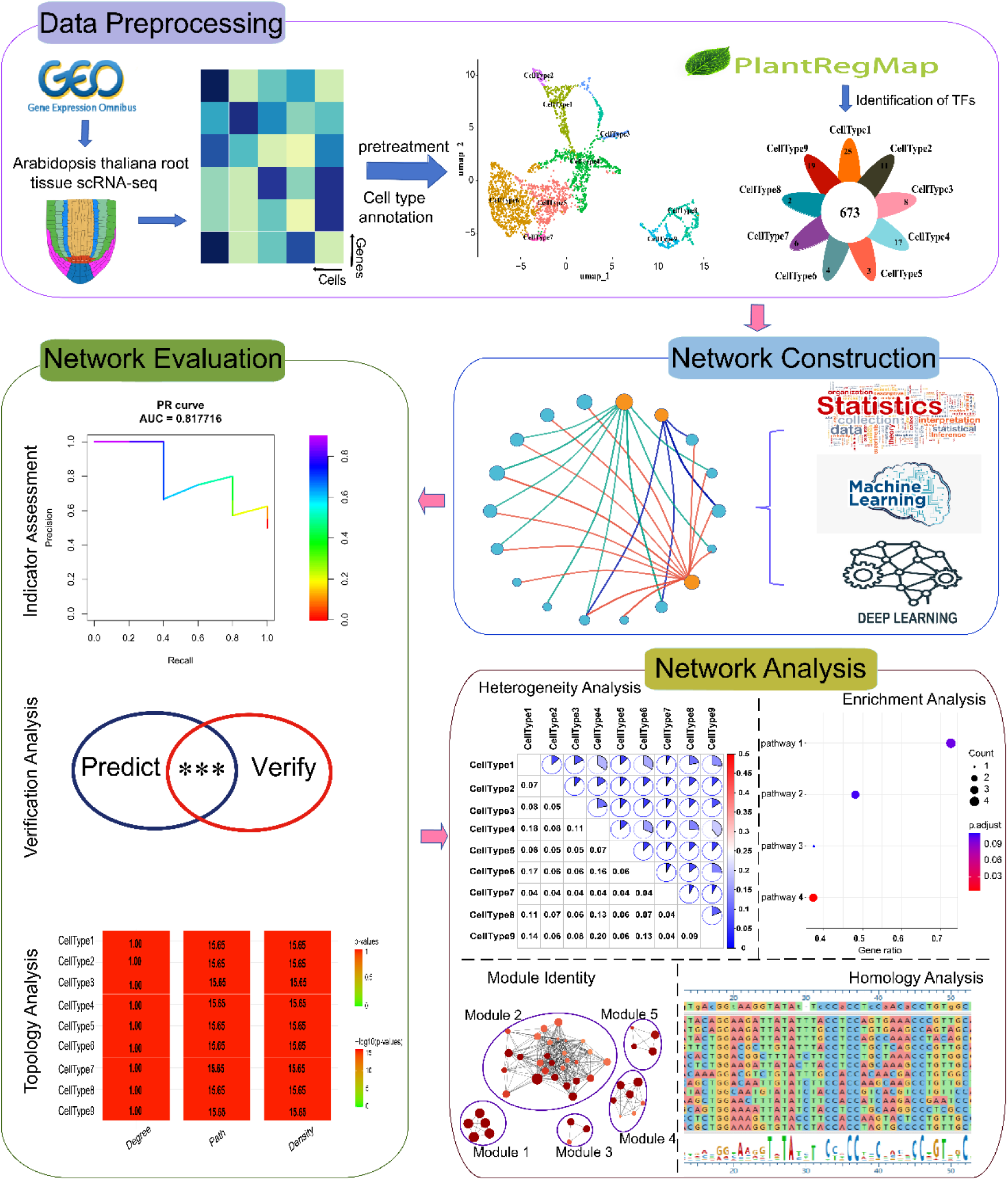
Research framework for constructing and analyzing cell-type-specific GRNs in *Arabidopsis thaliana* roots.

### Data download and data preprocessing

Single-cell sequencing data from *Arabidopsis* root (Accession ID: GSE123818) were obtained from the GEO database (https://www.ncbi.nlm.nih.gov/geo/)[37]. RNA-seq datasets from wild-type cell types were selected, and the raw expression matrices were transformed into Seurat objects using the CreateSeuratObject() function in R. Initial quality control(QC) filtering was performed to exclude low-quality cells and genes with low expression[38–40]. The proportion of mitochondrial genes was calculated, and further filtration of low-quality cells based on QC metrics. Data normalization was performed using the LogNormalize method, and the top 15 principal components were selected as the input parameters for UMAP dimensionality reduction based on principal component analysis (PCA) and the elbow method.

### Single-cell clustering analysis and annotation

We performed cell clustering using the FindClusters function and identified the cell types of the clusters. In order to annotate cells more accurately, we referenced known marker genes from the PlantCellMarker database (https://www.tobaccodb.org/pcmdb/homePage) to identify cell types, and further searched for differential marker genes among cell subsets for cell type annotation[41].

### Expression profiles analysis

The total count of genes for each cell type was measured, followed by calculating the overlap of genes between different cell types and counting TFs. These data were visualized using GraphPad Prism (version10.1.2). TF expression value from each cell type were averaged and z-transformed, resulting in a scaled expression values between −1.5 and +1.5 for each gene. Hierarchical Clustering was calculated by hclust() function in R, and the expression profiles were visualized as heatmaps generated by the pheatmap package (version1.0.12).

### Gene regulatory network inference

Three representative algorithms based on statistics, traditional machine learning and deep learning principles—WGCNA[42], GENIE3[43] and DeepSEM[44]—were employed to construct cell type-specific GRNs. We used the “WGCNA” R package to compute Pearson correlation between genes in each cell type’s expression matrix, filtering for significant co-expression relationships (*p*<0.05). GENIE3 employed random forest regression (parameters: K=“sqrt”, nCores=4, nTrees=1000) to assess regulatory weight of TFs on target genes. Edges were normalized using Z-score across the network, and only those with statistical significance (*p* < 0.05) were retained. DeepSEM integrated variational autoencoders (n_hidden=128, K=2, alpha=1, lr=1e-4) and graph convolutional networks, optimizing network structures through an alternating training strategy, ultimately outputting statistically significant regulatory edges (*p*<0.05) as the final network. The complete code is accessible at: GitHub (https://github.com/Jiangyu1220/ctsGRN).

### Analysis of network topology characteristics

The Kolmogorov-Smirnov (KS) test[45] was employed to evaluate whether the node degree distribution followed a power-law characteristic (*p* < 2.20×10^-16^), thereby determining if the network exhibited scale-free properties. The R package igraph was used to calculate topological parameters including node degree distribution, characteristic path length, and network density. Comparative analysis with random networks was performed using Student’s t-test[46] to assess whether the GRNs possessed small-world characteristics. All analyses were conducted based on a directed network model to ensure the results aligned with the actual features of biological regulatory networks.

### Network performance and quality assessment

The ChIP-seq data for TFs and open chromatin profiles (ATAC-seq, DNase-seq) from *Arabidopsis* roots were obtained from the ConnecTF database (https://ConnecTF.org) and ChIP-Hub database (https://biobigdata.nju.edu.cn/ChIPHub/) (S1 Table). TF binding sites located in accessible chromatin regions were kept and linked to their target genes for validating GRN quality. The performance of the algorithm was evaluated using the area under the precision-recall curve (AUPR) metric[47], computed with the PRROC package. The hypergeometric test is employed to evaluate whether the GRN exhibits predictive performance for TF-target gene regulatory relationships. The significance p-value of the test is calculated as follows:

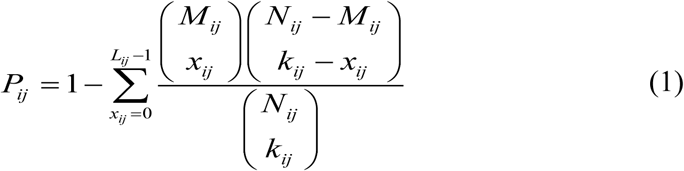

Where *N*_*ij*_ represents all possible regulatory relationships in the cell type, *M*_*ij*_ and *k*_*ij*_ denote the number of validated regulatory relationships among all possible relationships in the cell type and the number of predicted regulatory relationships validated in the cell type, respectively, and *L*_*ij*_ is the number of interactions between transcription factor *i* and target gene *j*.

### Screening and characterization of key transcription factors

TFs with degree centrality greater than 2,000 in each cell type were identified as key regulators[14]. The Sim.ceRNet function[48] was used to calculate the similarity among these key TFs across different cell types, revealing both commonalities and divergences between GRNs. For TFs shared among cell types, the mean (μ) and standard deviation (σ) of their degree centrality (number of regulatory edges) were calculated, and the coefficient of variation (CV = σ/μ) for these regulatory edges was ultimately computed.

### Yeast one-hybrid assay

Yeast one-hybrid (Y1H) assay was performed using Clontech’s Matchmaker Gold Yeast One-Hybrid (Y1H) Library Screening kit according to the manufacturer’s instructions. The full-length *AT1G62300* (*WRKY6*) was amplified and cloned into the pGADT7 vector to generate the construct AD-WRKY6. The promoter fragments of the *AT4G22710* (*CYP706A2*) gene containing the binding cis-elements were sequenced and introduced into the pAbAi vector by homologous recombination. The resultant yeast transformants were tested for self-activation of the CYP706A2 gene and used in the Y1H assays. Different combinations were co-transformed into the yeast strain Y1H Gold. The transformants were cultured in synthetic defined media (SD/-Leu) with or without 50 ng/ml of Aureobasidin A (AbA) for 4-5 days at 30 ℃, and the interactions were examined.

### Homology analysis

Protein sequences of Brassica napus orthologs were retrieved from BioMart (Ensembl Genomes 31) [49]. Multiple sequence alignment was performed using the ClustalW algorithm. A phylogenetic tree was constructed using MEGA11 software employing the neighbor-joining (NJ) method with 2,000 bootstrap replicates, and only branches with bootstrap support greater than 70% were retained[50].

### Functional module identification and enrichment analysis

GRNs were clustered into functional modules via the unsupervised Markov clustering algorithm (MCL) [51], Gene Ontology (GO) enrichment analysis of these functional modules was performed using the enrich GO() function from the R package clusterProfiler (v4.0.5), utilizing *A.thaliana* gene annotations from TAIR, p-values were calculated by Fisher’s exact test (pvalueCutoff = 0.05, pAdjustMethod = “BH”), and statistical significance was defined as a false discovery rate (FDR) < 0.05 after correcting for multiple testing.

## Declarations

## Ethics approval and consent to participate

The *Arabidopsis* data used in this study were obtained from publicly available databases. No permissions were necessary to use these data.

## Consent for publication

The authors of this paper all consent to its publication.

## Availability of data and materials

The data that support the findings of this study are available in the supporting information of this article. All data supporting the findings of the current study are listed in the ctsGRN framework (https://github.com/Jiangyu1220/ctsGRN)

## Authors’ contributions

Yu Jiang: Conceptualization, Data curation, Formal analysis, Investigation, Methodology, Software, Validation, Visualization, Writing - original draft. Haolin Yang: Formal analysis, Validation. Jian Gao: Formal analysis. Qi Zhang: Formal analysis. Zitai Yue: Formal analysis. Xuemei Wei: Conceptualization, Funding acquisition, Project administration, Resources, Supervision, Writing - review & editing. Junpeng Zhang: Conceptualization, Funding acquisition, Project administration, Resources, Supervision, Writing - review & editing.

## Acknowledgements

Not applicable.

