## Supplementary figures and images for "ctsGRN: Inferring cell type-specific gene regulatory networks in *Arabidopsis*"

### Supplementary Figure 1

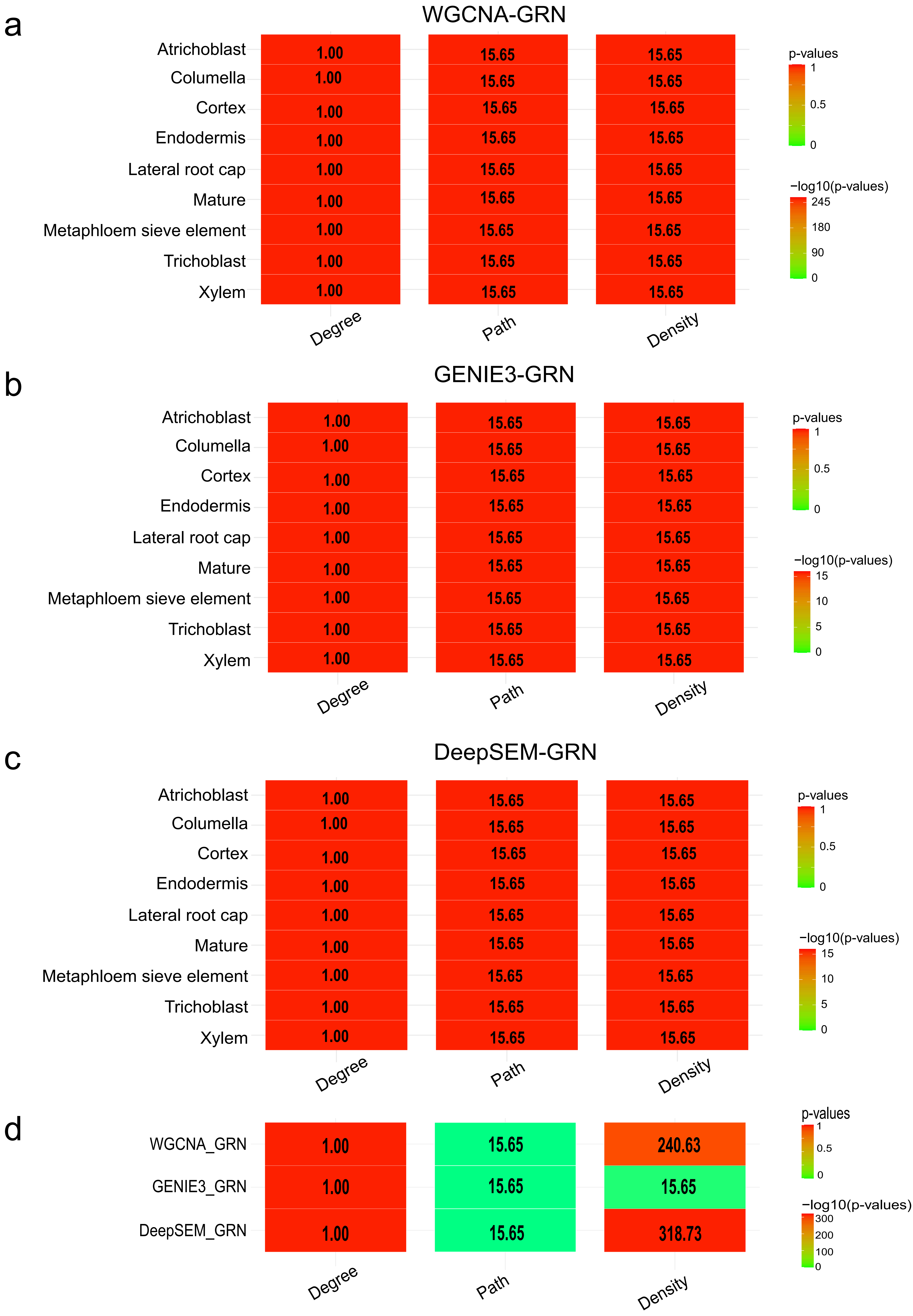

### Supplementary Figure 2

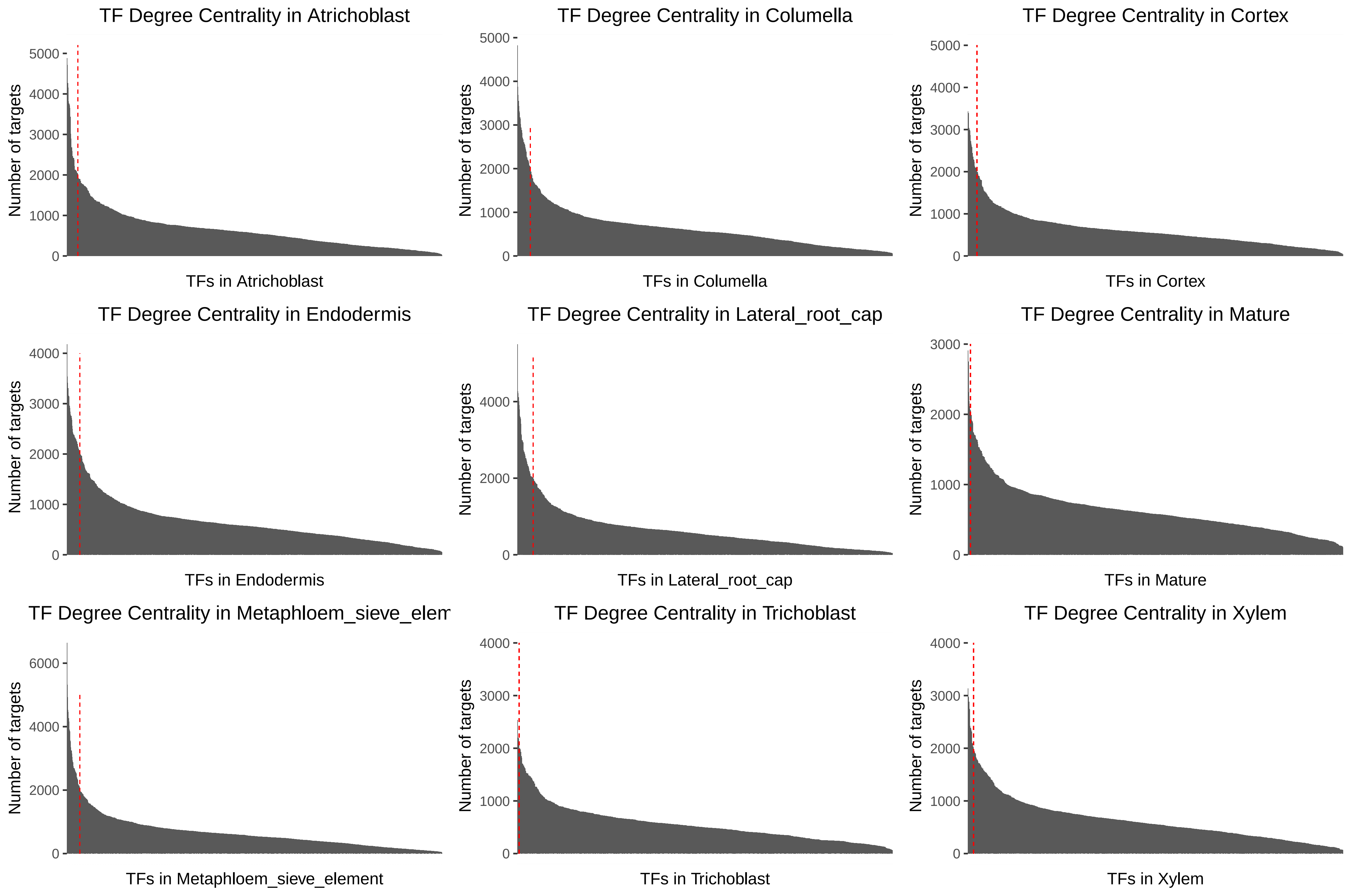

### Supplementary Figure 3

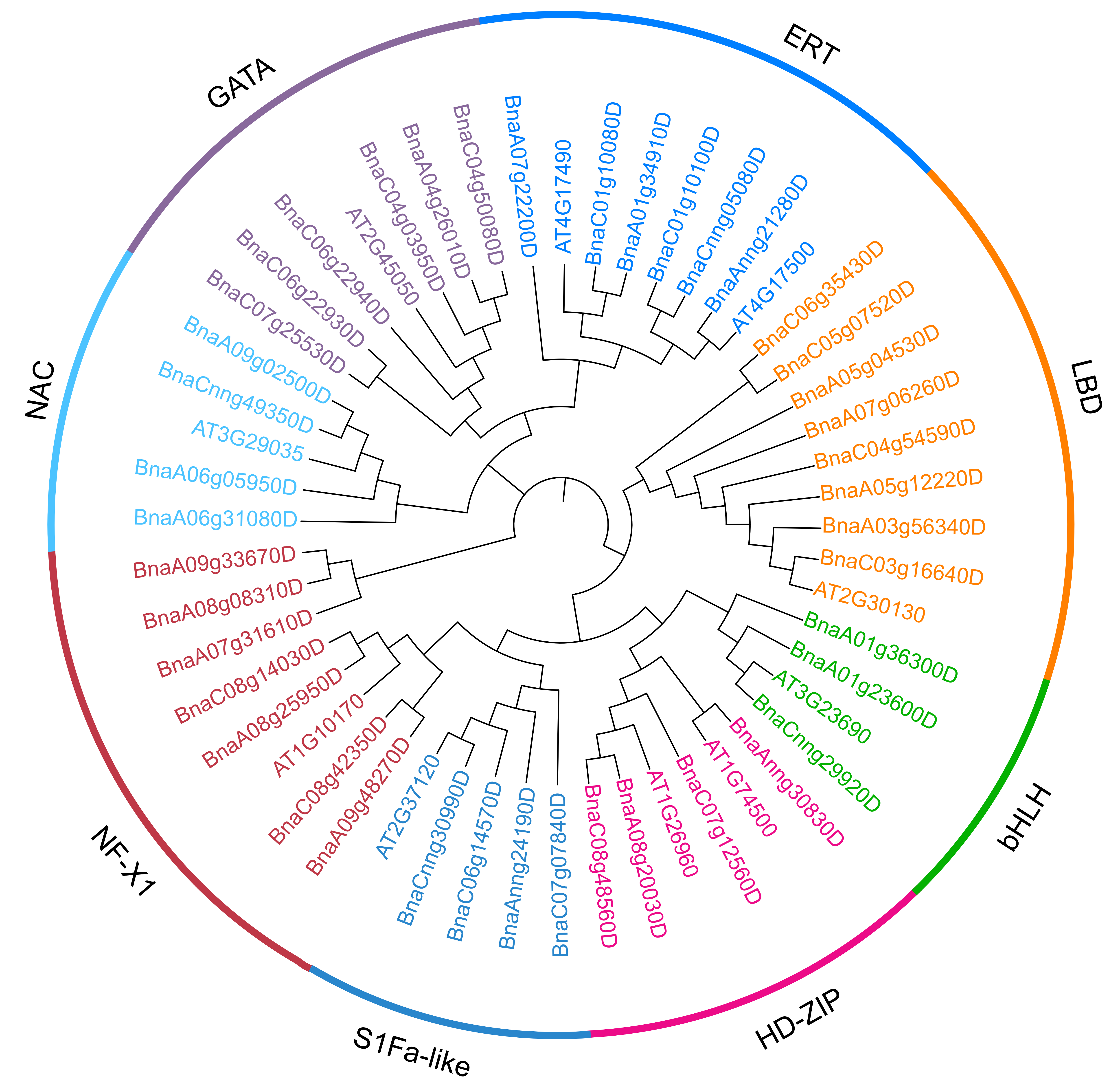
